# Enhanced nitric oxide synthesis through nitrate supply improves drought tolerance of sugarcane plants

**DOI:** 10.1101/860544

**Authors:** Maria D. Pissolato, Neidiquele M. Silveira, Paula J. Prataviera, Eduardo C. Machado, Amedea B. Seabra, Milena T. Pelegrino, Ladaslav Sodek, Rafael V. Ribeiro

## Abstract

Nitric oxide (NO) is an important signaling molecule associated with many biochemical and physiological processes in plants under stressful conditions. Nitrate reductase (NR) not only mediates the reduction of NO_3_^−^ to NO_2_^−^ but also reduces NO_2_^−^ to NO, a relevant pathway for NO production in higher plants. Herein, we hypothesized that sugarcane plants supplied with more NO_3_^−^ as a source of N would produce more NO under water deficit. Such NO would reduce oxidative damage and favor photosynthetic metabolism and growth under water limiting conditions. Sugarcane plants were grown in nutrient solution and received the same amount of nitrogen, with varying nitrate:ammonium ratios (100:0 and 70:30). Plants were then grown under well-watered or water deficit conditions, in which the osmotic potential of nutrient solution was −0.15 and −0.75 MPa, respectively. Under water deficit, plants exhibited higher root [NO_3_^−^] and [NO_2_^−^] when supplied with 100% NO_3_^−^. Accordingly, the same plants also showed higher root NR activity and root NO production. We also found higher photosynthetic rates and stomatal conductance in plants supplied with more NO_3_^−^, which improved root growth. ROS accumulation was reduced due to increases in the activity of catalase in leaves and superoxide dismutase and ascorbate peroxidase in roots of plants supplied with 100% NO_3_^−^ and facing water deficit. Such positive responses to water deficit were offset when a NO scavenger was supplied to the plants, thus confirming that increases in leaf gas exchange and plant growth were induced by NO. Concluding, NO_3_^−^ supply is an interesting strategy for alleviating the negative effects of water deficit on sugarcane plants, increasing drought tolerance through enhanced NO production. Our data also provide insights on how plant nutrition could improve crop tolerance against abiotic stresses, such as drought.

**Highlights:**

- Nitrate supply improves sugarcane growth under water deficit.
- Nitrate supply stimulated nitrate reductase activity and NO synthesis in sugarcane roots facing water deficit.
- Leaf gas exchange was increased by nitrate supply as well as root growth under water limiting conditions.
- Antioxidant responses were also improved in plants supplied exclusively with nitrate.
- Nitrogen management may be an interesting strategy for improving drought tolerance in sugarcane fields.

## Introduction

Nitric oxide (NO) is a diatomic radical gas and important signaling molecule in animals (Bogdan, 2015), fungi (Canovas *et al.*, 2016), bacteria (Crane *et al.*, 2010) and plants (Mur *et al.*, 2013). In plants, increasing evidence indicates NO as a key component of the signaling network, controlling numerous physiological and metabolic processes such as seed germination (Albertos *et al.*, 2015), flowering (He *et al.*, 2004), root growth (Fernandez-Marcos *et al.*, 2011), respiration, stomatal conductance (Moreau *et al.*, 2010; Wang *et al.*, 2015) and adaptive responses to biotic and abiotic stresses (Shan *et al.*, 2015; Fatma, *et al.*, 2016).

NO synthesis is increased in plants under drought and its role in promoting adaptive responses to cope with water deficit has been suggested (Cai *et al.*, 2015; Silveira *et al.*, 2017a). NO and NO-derived molecules play a critical role in intracellular redox signaling and in the activation of antioxidant defense mechanisms (Shi *et al.*, 2014; Hatamzadeh *et al.*, 2015; Silveira *et al.*, 2015). For example, NO supply conferred drought tolerance to wheat seedlings, reducing membrane damage (Garcia-Mata and Lamattina, 2001). Spraying S-nitrosogluthatione (GSNO) – a NO donor – on sugarcane plants resulted in higher photosynthesis under drought, promoting plant growth under stressful condition (Silveira *et al.*, 2016).

The protective action of exogenous NO donors has been attributed to the elimination of superoxide (O_2_^·-^) and enhancement of the antioxidant system in sugarcane plants under drought (Silveira *et al.*, 2017b). In addition, one of the main downstream effects of NO is the post-translational regulation involving thiols (Hancock and Neill, 2019). S-nitrosylation is a redox modification consisting in the reversible attachment of NO to the thiol group of a cysteine residue in a target protein leading to the formation S-nitrosothiols (SNOs) (Astier *et al.*, 2012; Fancy *et al.*, 2016). Then, S-nitrosylation may cause a conformational change in proteins, changing their activity or function. On the other hand, NO can react with reduced glutathione (GSH), producing S-nitrosoglutathione (GSNO) – an endogenous NO reservoir and an efficient NO donor (Jahnová *et al*., 2019).

While the mechanisms of NO synthesis in animals have been well documented, NO synthesis and its regulation in plants are complex and poorly understood. In animals, NO is bio-synthesized through NO synthase (NOS), which oxidizes *L*-arginine and produces *L*-citrulline and NO (Alderton *et al.*, 2001). Although some evidence indicates the presence of NOS-like activity in many plant species, genes encoding NOS have not yet been identified in higher plants (Hancock and Neill, 2014; Santolini *et al.*, 2017; Hancock and Neill, 2019). NO production in plant species and under diverse biological conditions point to the co-existence of multiple pathways, likely functioning in distinct tissues/organs and subcellular compartments (León and Costa-Broséta, 2019).

One of the most important pathways for NO production in land plants is through nitrate reductase (NR) (Gupta *et al.*, 2011; Fancy *et al.*, 2016; Chamizo-Ampudia *et al.*, 2017; León and Costa-Broséta, 2019), a multifunctional enzyme that catalyzes NO_3_^−^ reduction to NO_2_^−^, which is then reduced to NH_4_^+^ during the N assimilatory pathway (Heidari *et al.*, 2011). Arasimowicz-Jelonek *et al.* (2009) reported low NO concentration in cucumber seedlings treated with a NR inhibitor, suggesting its role in NO synthesis. In rice roots, NO production through NR was increased in response to NO_3_^−^ supply (Sun *et al.*, 2015). Furthermore, low NO production by *Physcomitrella patens* occurred when plants received a NR inhibitor (Andrés *et al.*, 2015). Although there are data supporting the association between NR activity and NO production in plants (Mur *et al.*, 2013), some authors have argued that NO production through NR represents only a small fraction (1-2%) of total NO_3_^−^ reduction (Yamasaki *et al.*, 1999; Rockel *et al.*, 2002). However, the role of such a NO production pathway and its sensitivity to small changes in NO_3_^−^ supply in plants under water deficit remain unknown.

Nitrogen is the most influential plant nutrient in sugarcane cultivation (Meyer *et al.*, 2007). Nitrate (NO_3_^−^), ammonium (NH_4_^+^), and urea (CO(NH_2_)_2_) are the main forms of fertilizers and, thus, are the main sources of N for crops (Esteban *et al.*, 2016). Some crops have a preference for NH_4_^+^ uptake (Malagoli *et al.*, 2000), but most studies have reported stress symptoms associated with NH_4_^+^ toxicity (Barreto *et al.*, 2018; Boschiero *et al.*, 2019). While Robinson *et al.* (2011) reported the sugarcane preference for NH_4_^+^, Pissolato *et al.* (2019) found that increasing NH_4_^+^ supply causes biomass reduction and photosynthesis impairment of sugarcane plants. Changing the N source, NO_3_^−^ supply has been shown to increase the tolerance to abiotic stresses in maize (Rios-Gonzalez *et al.*, 2002; Zhang *et al.*, 2012), wheat (Speer *et al.*, 1994), pea (Frechilla *et al.*, 2001), *Populus simonii* (Meng *et al.*, 2016) and grass species (Wang and Macko, 2011).

The literature concerning NO_3_^−^ supply and stress tolerance, taken together, led us to hypothesize that the increased plant performance under limiting conditions could be related to NO production through NR activity. Here, our aim was to test the hypothesis that sugarcane plants that receive NO_3_^−^ and no NH_4_^+^ as sources of nitrogen will have higher NR activity and thereby produce more NO, compared to plants receiving the same amount of nitrogen but as a mixture of NO_3_^−^ (70%) and NH_4_^+^ (30%). As a consequence of NO production, oxidative damage will be reduced under water deficit, favoring photosynthetic metabolism and plant growth.

## Materials and Methods

### Plant material and growth conditions

Pre-sprouted sugarcane seedlings *(Saccharum* spp.) cv. IACSP95-5000 developed by the Sugarcane Breeding Program of the Agronomic Institute (ProCana, IAC, Brazil) were used. Six-week-old plants were transferred to plastic boxes (4 L) containing nutrient solution modified from De Armas *et al.* (1992): 5 mmol L^−1^ N (nitrate 90% + ammonium 10%); 9 mmol L^−1^ Ca; 0.5 mmol L^−1^ Mg; 1.2 mmol L^−1^ P; 1.2 mmol L^−1^ S; 24 μmol L^−1^ B; 16 μmol L^−1^ Fe; 9 μmol L^−1^ Mn; 3.5 μmol L^−1^ Zn; 1 μmol L^−1^ Cu; and 0.1 μmol L^−1^ Mo. Plants received this solution for two weeks until the establishment of treatments and the nutrient solution was renewed every three days throughout the experimental period.

Electrical conductivity of nutrient solution was maintained between 1.8 and 2.0 mS cm^−1^ and pH at 5.9±0.1. The pH was adjusted daily with 0.5 M ascorbic acid or 0.5 M NaOH. Both variables were monitored on a daily basis using a portable electrical conductivity meter (mCA 150P, MS Tecnopon Instrumentação, Piracicaba SP, Brazil) and a portable pH meter (mPA 210P, MS Tecnopon Instrumentação, Piracicaba SP, Brazil), respectively. The nutrient solution volume was also checked daily and completed with water when necessary. The nutrient solution was aerated continuously by using an air compressor (Master Super II, Master, São Paulo SP, Brazil).

The experiment was carried in a growth chamber (Instalafrio, Brazil), with a 12 h photoperiod, air temperature of 30/20 °C (day/night), air relative humidity of 80% and photosynthetic photon flux density (PPFD) about 800 μmol m^−2^ s^−1^.

### Experiment I: Inducing NO production under water deficit through nitrate supply

Our previous study revealed that sugarcane plants can be supplied with 30% NH_4_^+^ in nutrient solution without compromising their photosynthesis and growth (Pissolato *et al.*, 2019). Thus, the NO_3_^−^:NH_4_^+^ ratios 100:0 and 70:30 were chosen to represent the treatments with more and less NO_3_^−^, while supplying the same amount of nitrogen and avoiding NH_4_^+^ toxicity. Plants were also subjected to varying water availability, according to the osmotic potential of nutrient solution: −0.15 MPa (reference, well-hydrated); and −0.75 MPa (water deficit, WD). The water deficit was induced by adding polyethylene glycol (Carbowax™ PEG-8000, Dow Chemical Comp, Midland MI, USA) to the nutrient solution, seven days after imposing NO_3_^−^:NH_4_^+^ ratios. To prevent osmotic shock, PEG-8000 was gradually added to the nutrient solution, reducing the osmotic potential of the solution by −0.20 MPa per day, i.e. −0.75 MPa was reached after three days (3^th^ day of the experiment). Plants were allowed to recover from water deficit after returning them to control conditions on the 7^th^ day of the experiment. They remained for 4 days under such conditions, when the experiment ended. For the biochemical analyses, leaf and root samplings were collected at the maximum water deficit (7^th^ day) and at the end of the recovery period (11^th^ day). Samples were collected, immediately immersed in liquid nitrogen and then stored at −80 °C.

### Leaf gas exchange

Gas exchange and chlorophyll fluorescence of the first fully expanded leaf with visible ligule were measured throughout the experimental period using an infrared gas analyzer (Li-6400, Licor, Lincoln NE, USA) equipped with a modulated fluorometer (6400-40 LCF, Licor, Lincoln NE, USA). Leaf CO_2_ assimilation (*A*_n_), stomatal conductance (*g*_s_) and the effective quantum efficiency of photosystem II (□_PSII_) were measured under PPFD of 2000 μmol m^−2^ s^−1^ and air CO_2_ concentration of 400 μmol mol^−1^. The measurements were performed between 10:30 and 12:30 h, as carried out previously (Pissolato *et al.*, 2019). The vapor pressure difference between leaf and air (VPDL) was 2.1±0.2 kPa and leaf temperature was 30±0.4°C during the evaluations.

### Chlorophyll content and leaf relative water content (RWC)

A chlorophyll meter (CFL 1030, Falker, Porto Alegre RS, Brazil) was used to assess the relative chlorophyll content (Chl). The relative water content was calculated using the fresh (FW), turgid (TW) and dry (DW) weights of leaf discs according to Jamaux *et al.* (1997): RWC=100×[(FW-DW)/(TW-DW)]. Measurements were taken at the maximum water deficit (7^th^ day), and four days after returning plants to the control condition (recovery period, 11^th^ day).

### Photosynthetic enzymes

The activity of ribulose-1,5-bisphosphate carboxylase/oxygenase (Rubisco, EC 4.1.1.39) was quantified in approximately 200 mg of leaves, which were macerated and homogenized in 100 mM bicine-NaOH buffer (pH 7.8), 1 mM ethylenediaminetetraacetic (EDTA), 5 mM MgCl_2_, 5 mM dithiothreitol (DTT), 1 mM phenylmethylsulfonyl fluoride (PMSF) and 10 μM leupeptin. The resulting solution was centrifuged at 14.000 *g* for 5 min at 4 °C. An aliquot of leaf extract was incubated with the reaction medium containing 100 mM bicine-NaOH (pH 8.0) 10 mM NaHCO_3_, 20 mM MgCl_2_, 3.5 mM ATP, 5 mM phosphocreatine, 0.25 mM NADH, 80 nkat glyceraldehyde-3-phosphate dehydrogenase, 80 nkat 3-phosphoglyceric phosphokinase and 80 nkat creatine phosphokinase, for 10 min at 25 °C. The oxidation of NADH was initiated by adding 0.5 mM ribulose-1,5-bisphosphate (RuBP) and total Rubisco activity was measured. The reduction of absorbance at 340 nm was monitored for 3 min (Sage *et al.*, 1988; Reid *et al.*, 1997).

The activity of phosphoenolpyruvate carboxylase (PEPC, EC 4.1.1.31) was also evaluated in approximately 200 mg of leaves, which were macerated and homogenized in 100 mM potassium phosphate buffer (pH 7), 1 mM EDTA, 1 mM PMSF and centrifuged at 14.000 *g* for 25 min at 4 °C. The supernatant was collected and the reaction medium for PEPC activity contained 50 mM Tris-HCl buffer (pH 7.8), 5 mM MgCl_2_, 5 mM glucose 6-phosphate, 10 mM NaHCO_3_, 33 nkat malic dehydrogenase and 0.3 mM NADH. The reaction was initiated by adding 4 mM phosphoenolpyruvate at 30 °C. The oxidation of NADH was monitored a 340 nm for 1 min (Degl’innocenti *et al.*, 2002).

Proteins were extracted from leaf samples with extraction buffer composed of 100 mM Tris, 1 mM EDTA, 5 mM DTT, 1 mM PMSF and separated by SDS-PAGE (Laemmli, 1970). The first gel was stained with Comassie Brilliant Blue and the second was used for Western blot. SDS-PAGE electrophoresis was performed with equal amounts of protein per lane. Soluble proteins were denatured using SDS and they were electrophoretically transferred to a nitrocellulose membrane (Towbin *et al.*, 1979). PEPC and Rubisco protein abundances were measured by detection of the PEPC subunit and Rubisco large subunit (RLS) using specific polyclonal antibodies (Agrisera Co, Sweden) according to the manufacturer’s instructions.

### Reactive oxygen species

The concentration of the superoxide anion (O_2_^·-^) was determined in 50 mg of fresh tissue incubated in an extraction medium consisting of 100 μM EDTA, 20 μM NADH, and 20 mM sodium phosphate buffer, pH 7.8. The reaction was initiated by adding 25.2 mM epinephrine in 0.1 N HCl. The samples were incubated at 28 °C under stirring for 5 min and the absorbance was read at 480 nm over a further 5 min (Mohammadi and Karr, 2001). O_2_^·-^ production was assessed by the accumulation of adrenochrome using a molar extinction coefficient of 4.0×10^3^ M^−1^ cm^−1^ (Boveris, 1984).

The quantification of hydrogen peroxide (H_2_O_2_) was performed following Alexieva *et al.* (2001). Homogenates were obtained from 100 mg of fresh tissue ground in liquid nitrogen with the addition of polyvinylpolypyrrolidone (PVPP) and 0.1% of trichloroacetic acid (TCA) solution (w/v). The extract was centrifuged at 10.000 *g* and 4 °C for 15 min. The reaction medium consisted of 1 mM KI, 0.1 M potassium phosphate buffer (pH 7.5) and crude extract. The microtubes were left on ice in the dark for 1 h. After this period, the absorbance was read at 390 nm. A standard curve was obtained with H_2_O_2_ and the results were expressed as μmol H_2_O_2_ g^−1^ FW.

### Lipid peroxidation

The concentration of malondialdehyde (MDA) was measured and used as a proxy of lipid peroxidation. 200 mg of fresh tissue were macerated in extraction medium containing 0.1% TCA (w/v) and centrifuged at 10.000 *g* for 15 min. The supernatant was added to 0.5% thiobarbituric acid (w/v) in 20% TCA (w/v), and the mixture incubated at 95 °C for 20 min (Cakmak and Horst, 1991). After this time, the reaction was stopped in an ice bath. Then a new centrifugation was performed at 10.000 *g* for 10 min, and after 30 min at room temperature the absorbance was read at 532 and 600 nm and the non-specific absorbance at 600 nm was discounted. The MDA concentration was calculated using an extinction coefficient of 155 mM^−1^ cm^−1^ (Heath and Packer, 1968) and results were expressed as nmol MDA g^−1^ FW

### Antioxidant activity and protein extraction

The crude enzymatic extracts for the determination of superoxide dismutase activity (SOD), catalase (CAT) and ascorbate peroxidase (APX) were obtained from 100 mg of plant tissue in specific medium, followed by centrifugation at 12.000 *g* for 15 min at 4 °C. The specific medium for CAT and SOD consisted of 0.1 M potassium phosphate buffer (pH 6.8), 0.1 mM EDTA, 1 mM PMSF and 1% PVPP, according to Peixoto *et al.* (1999). The specific medium for APX was composed of 50 mM potassium phosphate buffer (pH 7.0), 1 mM ascorbic acid and 1 mM EDTA (Nakano and Asada, 1981).

Superoxide dismutase (SOD, EC 1.15.1.1) activity was determined according to Giannopolitis and Ries (1977). The crude extract was added to the reaction medium consisting of 100 mM sodium phosphate buffer (pH 7.8), 50 mM methionine, 5 mM EDTA, deionized water, 100 μM riboflavin and 1 mM nitro blue tetrazolium chloride (NBT). A group of tubes was exposed to light (fluorescent lamp, 30 W) for 10 min, and another group remained in darkness. The absorbance was measured at 560 nm and one unit of SOD defined as the amount of enzyme required to inhibit NBT photoreduction by 50%, and activity expressed as U min^−1^ mg^−1^ of protein.

Catalase (CAT, EC 1.11.1.6) activity was quantified following the procedure described by Havir and McHale (1987). The crude extract was added to the reaction medium consisting of 100 mM potassium phosphate buffer (pH 6.8), deionized water and 125 mM H_2_O_2_. The reaction was carried out in a water bath at 25 °C for 2 min and CAT activity was assessed by the decrease in absorbance at 240 nm, using the molar extinction coefficient of 36 M^−1^ cm^−1^ and expressed activity as nmol min^−1^ mg^−1^ of protein.

Ascorbate peroxidase (APX, EC 1.11.1.11) activity was evaluated as described by Nakano and Asada (1981). The crude extract was added in reaction medium consisting of 100 mM potassium phosphate buffer (pH 6.8), deionized water, 10 mM ascorbic acid and 10 mM H_2_O_2_. The reaction was carried out at 25 °C for 2 min and APX activity quantified by the decrease in absorbance at 290 nm, using the molar extinction coefficient of 2.8 M^−1^ cm^−1^ and expressing activity as μmol min^−1^ mg^−1^ of protein.

The protein levels were determined by the Bradford method (Bradford, 1976), using bovine serum albumin (BSA) as the standard. The extract used for this analysis was the same as for SOD and CAT enzymes.

### Nitrate, nitrite and ammonium

Fresh plant tissue samples (500 mg) were ground in liquid nitrogen and extraction medium containing methanol:chloroform:water (12:5:3 v/v). After centrifugation at 2.000 *g* for 5 min, the supernatants were collected and chloroform and deionized water were added to them. The mixture was shaken vigorously and then centrifuged for 3 min at 2.000 *g* for phase separation. The upper aqueous phase was collected and maintained in a water bath at 37 °C to remove traces of chloroform and then the extracts were stored at −20 °C (Bieleski and Turner, 1966).

For nitrate determination, an aliquot of the extract was pipetted into test tubes containing reaction medium (5% salicylic acid in conc. H_2_SO_4_). After 20 min, 2 N NaOH was added and the solution stirred. After cooling to room temperature, the absorbance was read in a spectrophotometer at 410 nm and the nitrate content calculated from a standard curve using KNO_3_ (100-1000 nmol) (Cataldo *et al.*, 1975). For nitrite, an aliquot of the extract was added to 1% sulfanilamide solution in 3 N HCl and 0.02% N-naphthyl ethylenediamine solution. The tubes were allowed to stand for 30 min in the dark at room temperature. Deionized water was added and nitrite content quantified after reading the absorbance at 540 nm (Hageman and Reed, 1980). For ammonium, the extract was added to microtubes, where solution A (1% phenol and 0.005% sodium nitroprusside) was added and followed by solution B (0.5% sodium hydroxide containing 2.52% sodium hypochlorite). The tubes were incubated for 35 min in a water bath at 37 °C and the absorbance read at 625 nm after cooling to room temperature (McCullough, 1967). A standard curve of (NH_4_)_2_SO_4_ was used to estimate the ammonium content.

### Nitrate reductase (NR) activity

Leaf and root nitrate reductase (NR, EC 1.7.1.1) activity was estimated as the rate of nitrite (NO_2_^−^) production (Cambraia *et al.*, 1989). The enzyme extract was obtained from the macerate of 200 mg of fresh tissue with liquid nitrogen and homogenized with extraction medium containing 0.1 M tris-HCl buffer (pH 8.1), 4 mM NiSO_4_, 20 mM reduced glutathione (GSH), deionized water and 0.5 mM PMSF. Then, the crude extracts were centrifuged at 10.000 *g* for 10 min at 4 °C and the supernatant was collected and maintained on ice. The extract was added to the assay medium containing 100 mM Tris-HCl buffer (pH 7.5), 10 mM KNO_3_, 0.05 mM NADH and triton 1% X-100 (v/v), mixed and incubated at 30 °C for 10 min. The reaction was quenched by adding 1% sulfanilamide solution in 1 M HCl and 0.01% N-naphthyl ethylenediamine. Nitrite production was determined by absorbance at 540 nm using a standard curve with KNO_2_. The NR activity was expressed as nmol NO_2_^−^ min^−1^ mg^−1^ protein.

### S-nitrosogluthatione reductase (GSNOR) activity

Leaf and root S-nitrosogluthatione reductase (GSNOR, EC 1.2.1.1) activity was determined spectrophotometrically at 25 °C by monitoring the oxidation of NADH at 340 nm, based on Rodríguez-Ruiz *et al.* (2017). Briefly, 200 mg of fresh tissue were grounded with liquid nitrogen, resuspended in 20 mM HEPES buffer (pH 8.0), 10 mM EDTA, 0.5 mM PMSF and centrifuged for 10 min at 10.000 *g* and 4 °C. The enzyme extract was added in to the assay medium (20 mM HEPES buffer pH 8.0 and 1.8 mM NADH) at 25 °C, and maintained in the dark. The reaction was started by adding 4 mM GSNO (Silveira *et al.*, 2016) and the GSNOR activity followed by NADH oxidation at 340 nm. Activity was calculated using the NADH extinction coefficient (6.22 mM^−1^ cm^−1^ at 340 nm) and expressed as nmol NADH min^−1^ mg^−1^ protein.

### S-nitrosothiols content

The total leaf and root proteins were extracted in deionized water and the resulting homogenate was used to estimate the S-nitrosothiol content through an amperometer, as described by Santos *et al*. (2016) and Zhang *et al.* (2000). Measurements were performed with the WPI amperometer TBR 4100/1025 (World Precision Instruments Inc., Sarasota FL, USA) and a specific nitric oxide (NO) sensor, ISO-NOP (2 mm). Aliquots of aqueous suspension were added to the sample compartment containing aqueous copper chloride solution (0.1 mol L^−1^). This condition allowed the detection of free NO released from the S-nitrosothiols present in the leaf and root protein homogenate. The samples were run in triplicate and the calibration curve was obtained with newly prepared GSNO solutions. The data were compared with the standard curve obtained and normalized against fresh weight. The SNO content was expressed as μmol NO g^−1^ FW.

### Intracellular NO detection

NO was assayed in leaf and root segments. For the roots, it was collected approximately 1 cm from the middle part of secondary root. For the leaves, a thin cross section was made with the aid of a scalpel. The segments were incubated in MES-KCl buffer (10 mM MES, 50 mM KCl, 0.1 mM CaCl_2_, pH 6.15), at room temperature for 15 min. Then, these segments were incubated in solution of 10 μM DAF2-DA, mixing for 40 min in the dark at room temperature (Desikan *et al.*, 2002; Bright *et al.*, 2009). The samples were washed with buffer to remove the excess of DAF2-DA, placed onto a glass slide and covered with a glass slip before observing fluorescence using an inverted confocal microscope set for excitation at 488 nm and emission at 515 nm (Model Zeiss LSM510, Carl Zeiss AG, Germany). Photographs were taken with a 10x magnification, 15 s exposure and 1x gain. Images were analyzed using ImageJ software (NIH, Bethesda, MD, USA) and data were normalized by subtracting the values of the negative control (plants well-hydrated) and presented as mean pixel intensities.

### Biometry

Leaf and root dry masses were quantified after drying samples in an oven (60 °C) with forced-air circulation until constant weight. Leaf area of each plant was evaluated with a portable leaf area meter (model LI-3000, Li-Cor Inc., Lincoln NE, USA).

### Experiment II: Using cPTIO to offset the benefits of NO in plants under water deficit

An additional experiment was performed to verify whether the benefits found in plants supplied with only NO_3_^−^ and subjected to water deficit were in fact caused by NO. We used a NO scavenger, 2-(4-carboxyphenyl)-4,4,5,5-tetramethylimidazoline-1-oxyl-3-oxide (cPTIO). cPTIO is a stable organic radical developed by Akaike and Maeda (1996), which has been widely used as a control as it oxidizes the NO molecule to form NO_2_. In plants supplied with only NO_3_^−^ as N source, the following treatments were evaluated: (a) well-watered condition, with an osmotic potential of the nutrient solution of −0.15 MPa; (b) water deficit, with an osmotic potential of nutrient solution of −0.75 MPa; and (c) same as b with 100 μM cPTIO.

First, plants were moved and roots placed in a moist chamber, where they were sprayed with cPTIO and remained in the dark for 1 hour. After this treatment, the plants were returned to the boxes with the original nutrient solution. This procedure was performed for four consecutive days from the moment the water deficit (−0.75 MPa) was installed. We also evaluated the production of intracellular NO, plant biomass, leaf CO_2_ assimilation (*A*_n_) and stomatal conductance (*g*_s_) as described previously.

### Experimental design and statistical analyses

The experimental design was completely randomized and two causes of variation were analyzed: water availability and nitrogen source. Data were then subjected to an analysis of variance (ANOVA) and when statistical significance was detected, the mean values (*n*=4) were compared by the Tukey test (*p*<0.05) using the software Assistat version 7.7 (UFCG, Campina Grande PB, Brazil).

## Results

### Experiment I: Sugarcane responses to water deficit as affected by NO_3_^−^ supply

#### Relative water content and photosynthesis

A significant reduction in leaf relative water content was found under water deficit, as compared to well-watered conditions (Fig. 1d). The relative chlorophyll content was also reduced at the maximum water deficit, with no differences induced by NO_3_^−^ supply (data not shown). Low water availability also caused a large reduction in leaf CO_2_ assimilation (*A*_n_); however, plants supplied with more NO_3_^−^ exhibited higher photosynthetic rates than those under NO_3_^−^:NH_4_^+^ 70:30 (Fig. 1a). In addition, those plants showed a faster recovery of *A*_n_ when compared to ones receiving 70% NO_3_^−^ (Fig. 1a). Similar results were found for stomatal conductance (Fig. 1b) and effective quantum efficiency of PSII (Fig. 1b,c). We did not observe any significant difference among treatments for the PEPC abundance and activity at maximum water deficit (Suppl. Fig. S1a,c). However, both Rubisco abundance and activity were decreased under water deficit, regardless of the variation in NO_3_^−^ supply (Suppl. Fig. S1b,d).

**Figure 1.**
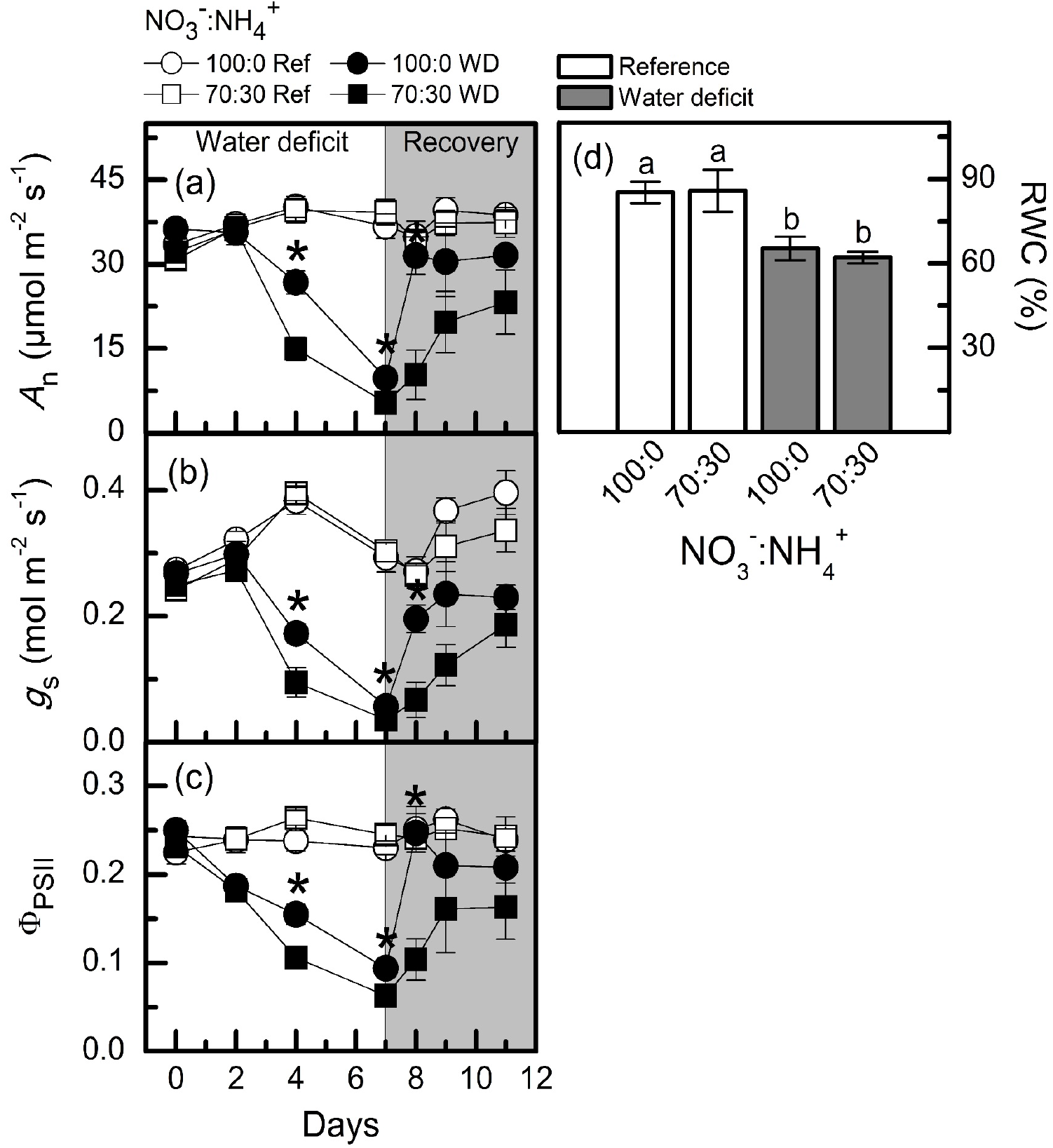
Leaf CO_2_ assimilation (*A*_n_, in a), stomatal conductance (*g*_S_, in b), effective quantum efficiency of PSII (Φ_PSII_, in c) and leaf relative water content (RWC, in d) in sugarcane plants maintained well-hydrated (ref, white symbols and bars) or subjected to water deficit (WD, black symbols and gray bars) and supplied with varying NO_3_^−^:NH_4_^+^ ratios: 100:0 and 70:30. The white area indicates the period of water deficit and the shaded area indicates the period of recovery. Symbols and bars represent the mean value of four replications ± se. Asterisks indicate significant differences between treatments under water deficit and different letters indicate statistical difference among treatments (Tukey test, *p*<0.05).

#### Nitrate and ammonium

Leaf [NO_3_^−^] was higher in water-stressed plants as compared to well-hydrated ones, but no difference was found due to NO_3_^−^ supply under low water availability (Fig. 2a). Root [NO_3_^−^] was significantly higher in plants supplied with 100% NO_3_^−^ and subjected to water deficit (Fig. 2b). While leaf [NO_2_^−^] did not vary among treatments (Fig. 2c), we found the highest root [NO_2_^−^] in plants supplied with 100% NO_3_^−^ under water deficit (Fig. 2d). We did not find significant changes in leaf and root [NH_4_^+^] due to NO_3_^−^ supply, regardless the water regime (Fig. 2e,f). During the recovery period, both previously stressed plants and the controls presented similar leaf and root [NO_3_^−^], [NO_2_^−^] and [NH_4_^+^] (Fig. 2).

**Figure 2.**
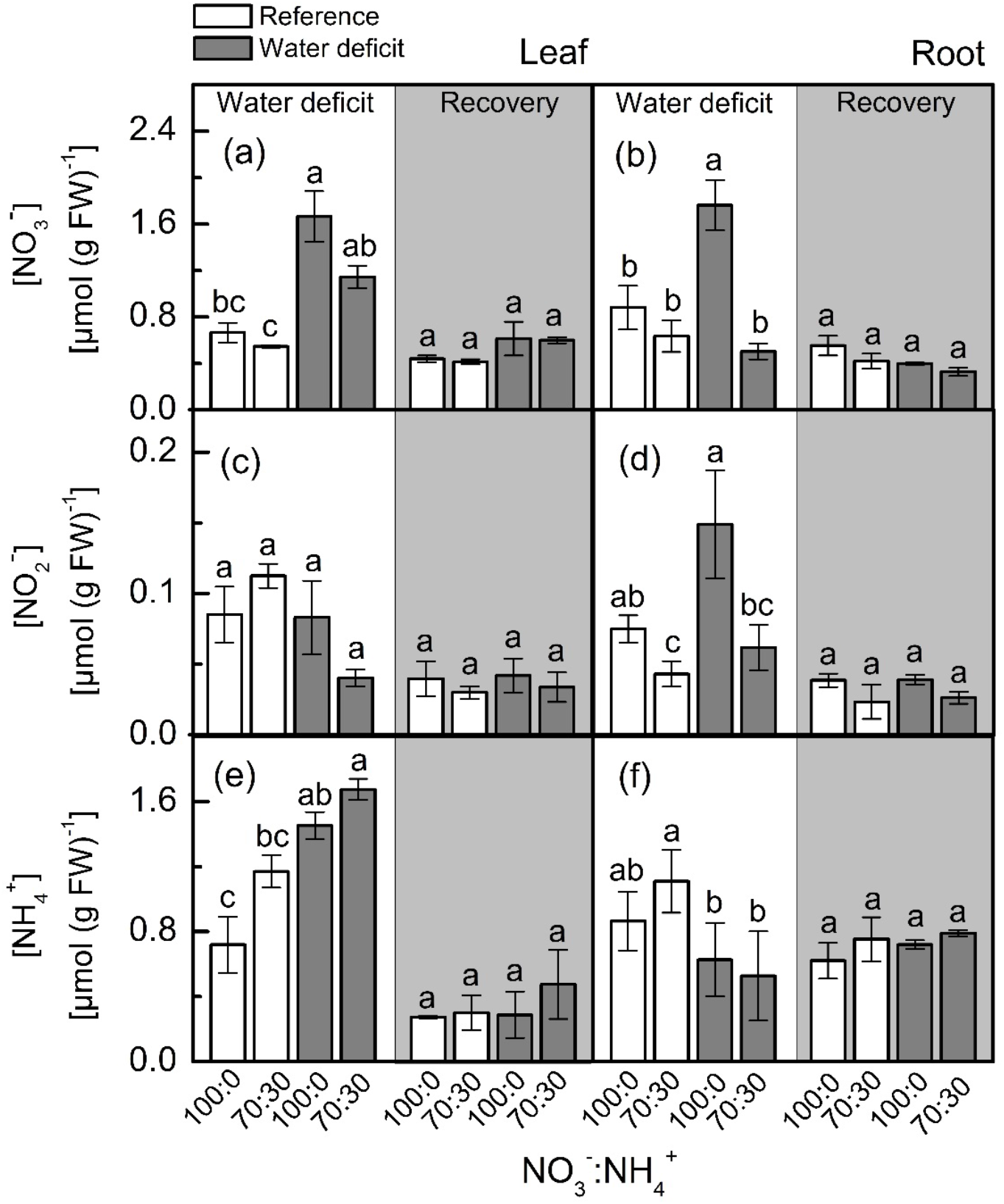
Concentration of nitrate (a and b), nitrite (c and d) and ammonium (e and f) in leaves (a, c and e) and roots (b, d and f) of sugarcane plants maintained well-hydrated (reference, white bars) or subjected to water deficit (gray bars) and supplied with varying NO_3_^−^:NH_4_^+^ ratios: 100:0 and 70:30. The white area indicates the period of water deficit and the shaded area indicates the period of recovery. Bars represent the mean value of four replications ± se. Different letters indicate statistical difference among treatments (Tukey test, *p*<0.05).

#### Nitrate reductase, S-nitrosoglutathione reductase and S-nitrosothiols

Under low water availability, nitrate reductase (NR) activity was higher in plants supplied with 100% NO_3_^−^ than those receiving 70% NO_3_^−^, regardless the plant organ (Fig. 3a,b). While we did not notice differences among treatments for leaf NR activity during the recovery period, root NR activity was higher under water deficit (Fig. 3b). Under water deficit, plants supplied with 100% NO_3_^−^ showed higher root GSNOR activity than those under 70% NO_3_^−^ (Fig. 3d). Nonsignificant differences were found in leaf SNO concentration while varying NO_3_^−^ supply (Fig. 3e). However, the lowest root S-nitrosothiols (SNO) concentration was observed in plants supplied with 100% NO_3_^−^ under water deficit (Fig. 3f).

**Figure 3.**
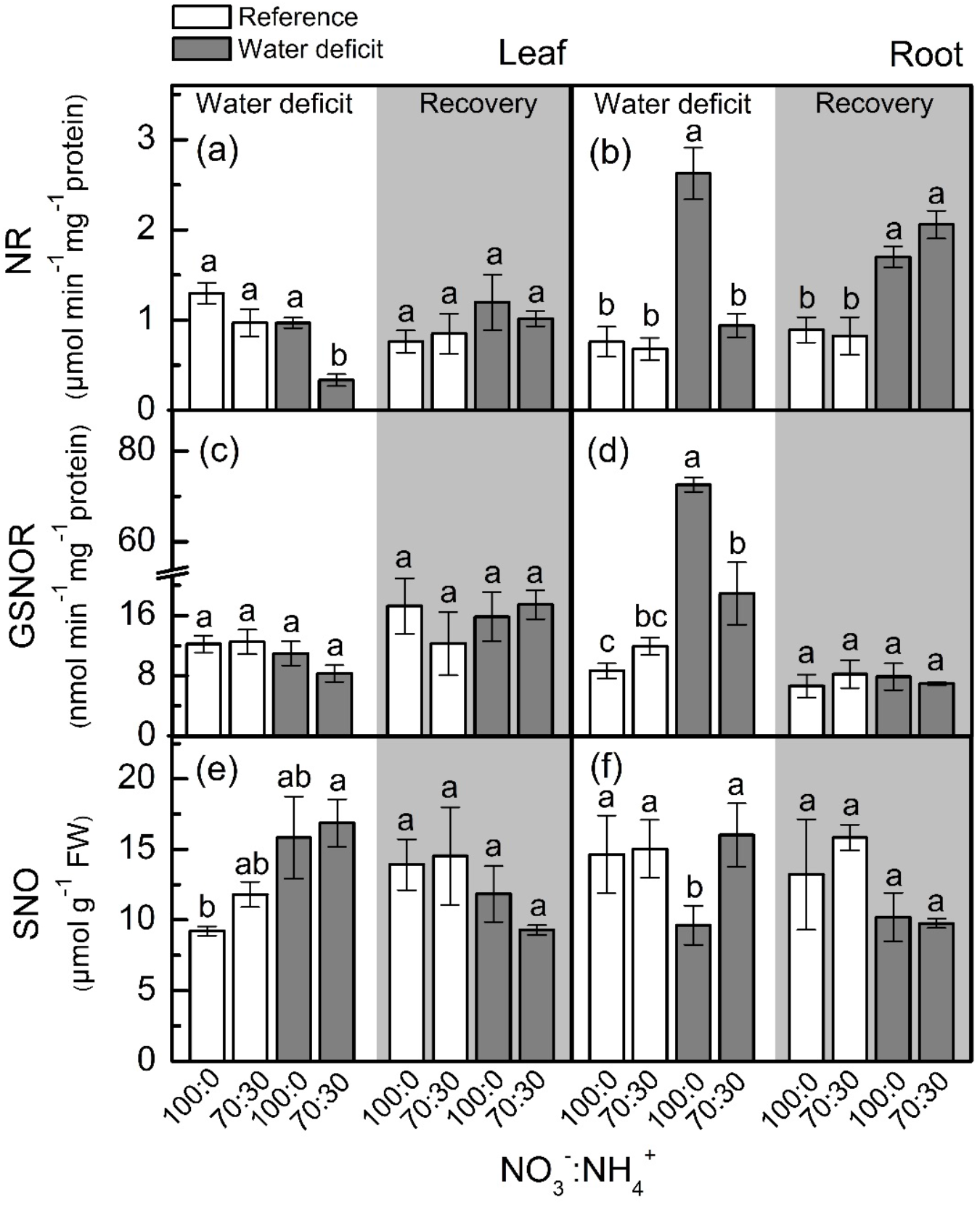
Nitrate reductase activity (NR, in a and b), S-nitrosoglutathione reductase activity (GSNOR, in c and d) and S-nitrosothiol concentration (SNO, in e and f) in leaves (a, c and e) and roots (b, d and f) of sugarcane plants maintained well-hydrated (reference, white bars) or subjected to water deficit (gray bars) and supplied with varying NO_3_^−^:NH_4_^+^ ratios: 100:0 and 70:30. The white area indicates the period of water deficit and the shaded area indicates the period of recovery. Bars represent the mean value of four replications ± se. Different letters indicate statistical difference among treatments (Tukey test, *p*<0.05).

#### Antioxidant metabolism

Plants supplied with less NO_3_^−^ presented higher leaf [O_2_^·-^] when compared to ones supplied with 100% NO_3_^−^ under water deficit (Fig. 4a). When plants faced water deficit, the highest root [H_2_O_2_] was found under 70% NO_3_^−^ supply (Fig. 4d). Although showing higher accumulation of O_2_^·-^ and H_2_O_2_ in leaves and roots, plants supplied with 70% NO_3_^−^ did not show higher MDA content than those under 100% NO_3_^−^ (Fig. 4e,f).

**Figure 4.**
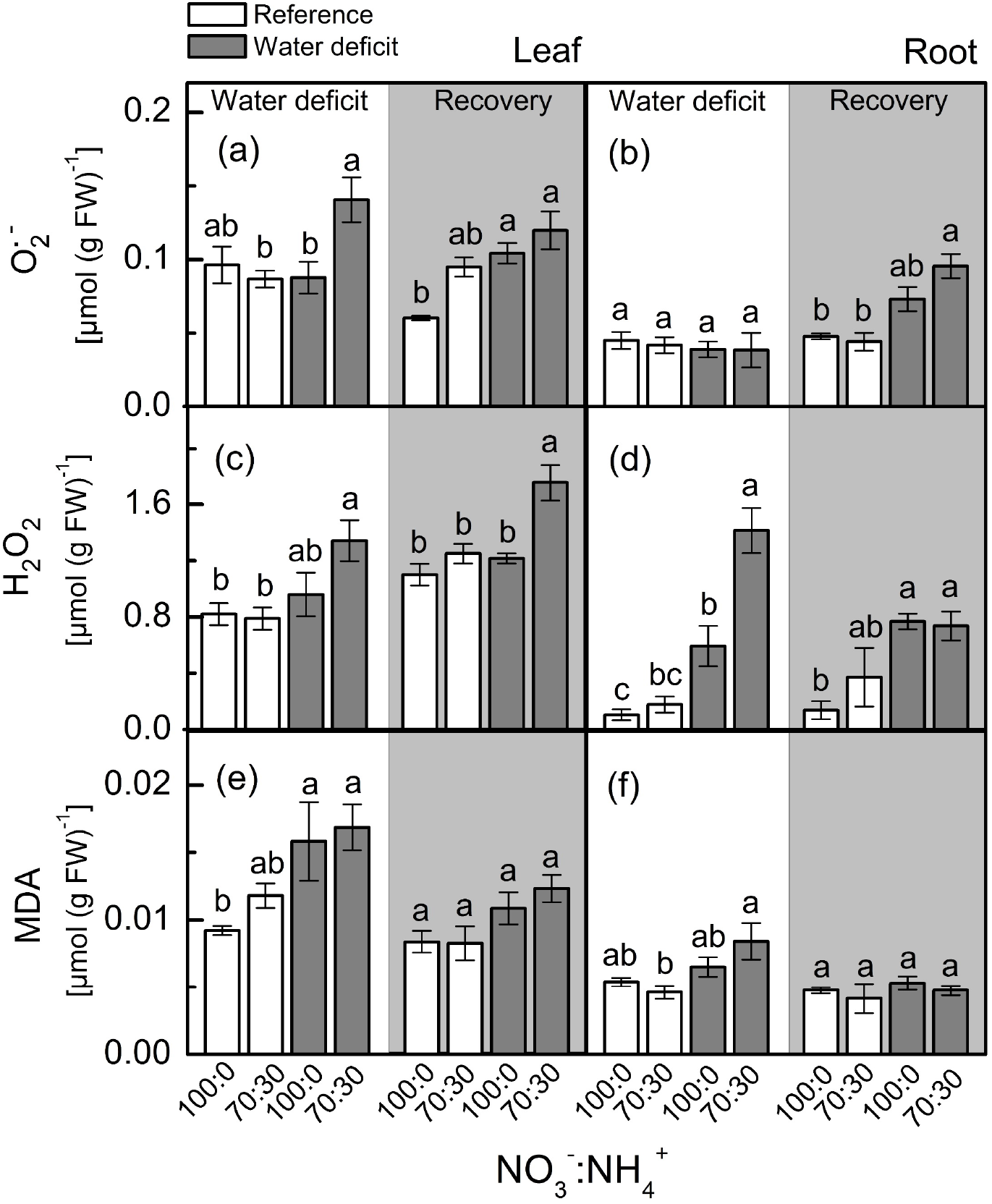
Concentration of superoxide anion (O_2_^·−^, a and b), hydrogen peroxide (H_2_O_2_, c and d) and malondialdehyde (MDA, in e and f) in leaves (a, c and e) and roots (b, d and f) of sugarcane plants maintained well-hydrated (reference, white bars) or subjected to water deficit (gray bars) and supplied with varying NO_3_^−^:NH_4_^+^ ratios: 100:0 and 70:30. The white area indicates the period of water deficit and the shaded area indicates the period of recovery. Bars represent the mean value of four replications ± se. Different letters indicate statistical difference among treatments (Tukey test, *p*<0.05).

At the maximum water deficit, the highest superoxide dismutase (SOD) activity was observed in roots supplied with 100% NO_3_^−^ (Fig. 5b), with no differences in leaf SOD activity due to changes in NO_3_^−^ supply (Fig. 5a). Root catalase activity was not changed by NO_3_^−^ supply and water deficit (Fig. 5f), but plants supplied with 100% NO_3_^−^ showed higher leaf catalase and root ascorbate peroxidase activities under water deficit (Fig. 5e,d).

**Figure 5.**
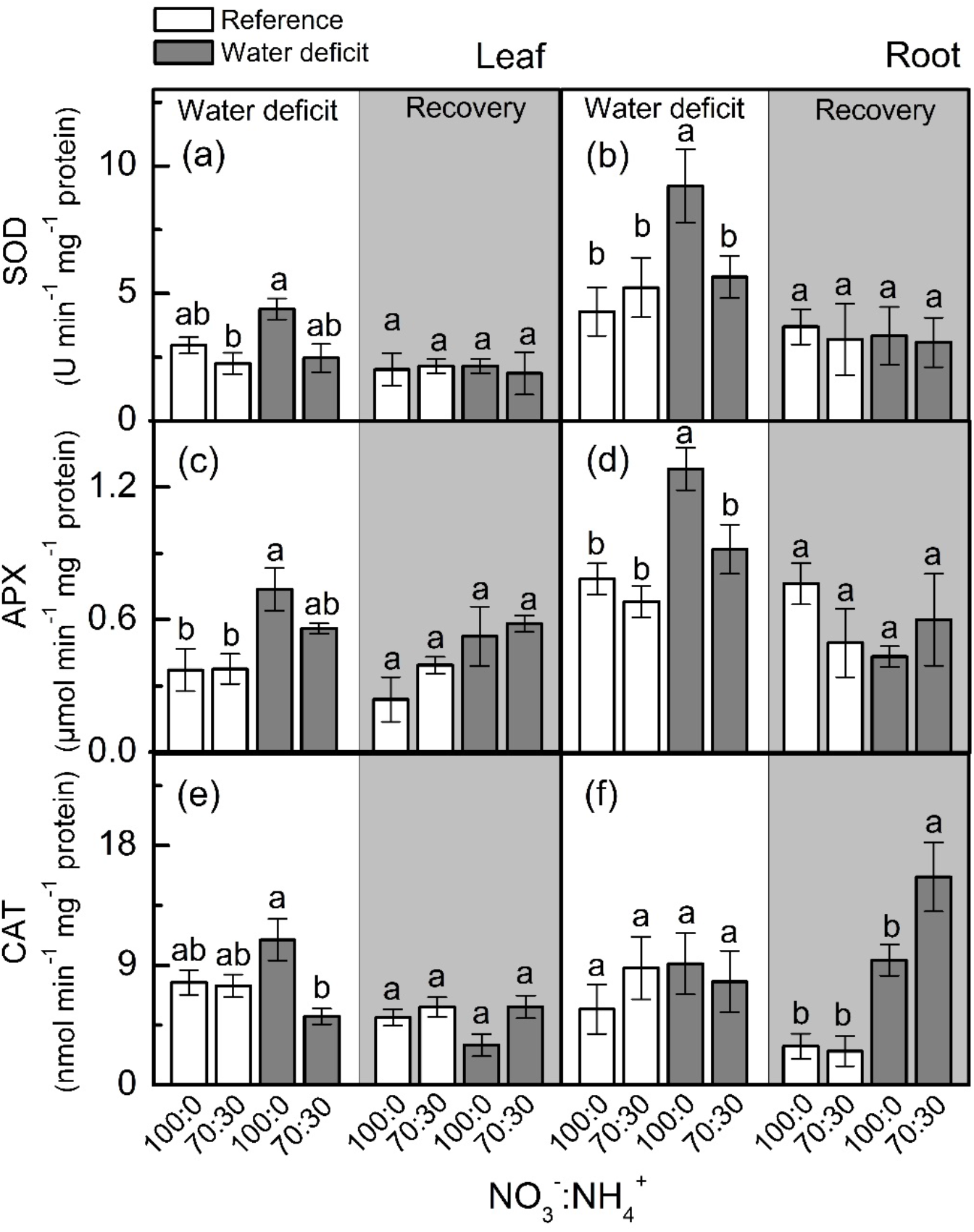
Superoxide dismutase activity (SOD, in a and b), ascorbate peroxidase activity (APX, in c and d) and catalase activity (CAT, in e and f) in leaves (a, c and e) and roots (b, d and f) of sugarcane plants maintained well-hydrated (reference, white bars) or subjected to water deficit (gray bars) and supplied with varying NO_3_^−^:NH_4_^+^ ratios: 100:0 and 70:30. The white area indicates the period of water deficit and the shaded area indicates the period of recovery. Bars represent the mean value of four replications ± se. Different letters indicate statistical difference among treatments (Tukey test, *p*<0.05).

#### Intracellular NO synthesis

When plants were facing low water availability, the intracellular NO was increased in both leaves and roots (Fig. 6). However, roots receiving 100% NO_3_^−^ exhibited higher NO production than those supplied with 70% NO_3_^−^ (Fig. 6b). Such a response did not occur in leaves (Fig. 6a).

**Figure 6.**
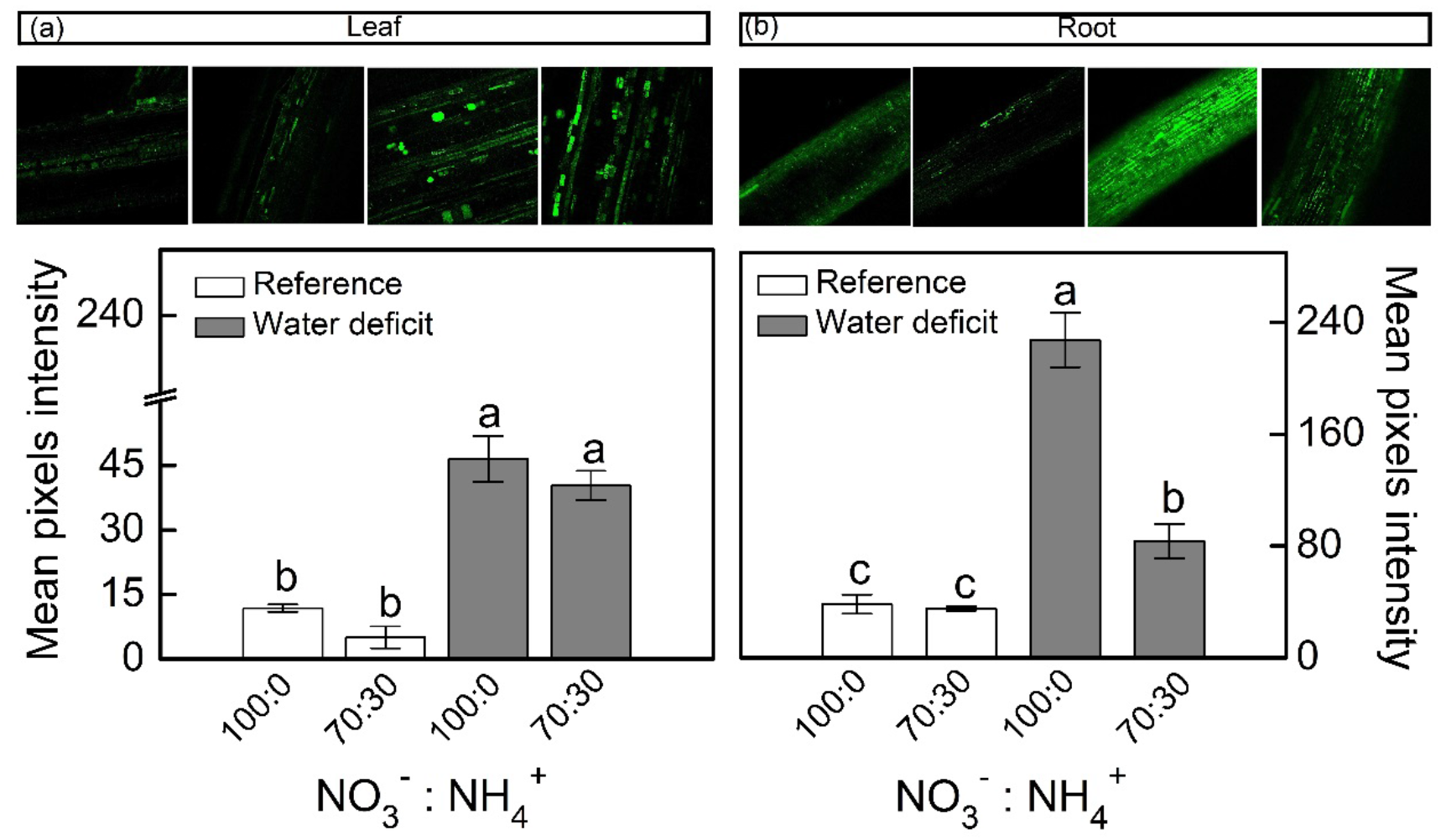
Confocal microscopy images showing intracellular NO synthesis in leaves (a) and roots (b) of sugarcane plants maintained well-hydrated (reference, white bars) or subjected to water deficit (gray bars) and supplied with varying NO_3_^−^:NH_4_^+^ ratios: 100:0 and 70:30. Mean pixel intensities are also shown. Bars represent the mean value of four replications ± se. Different letters indicate statistical difference among treatments (Tukey test, *p*<0.05).

#### Plant growth

The root dry mass of plants supplied with 70% NO_3_^−^ was significantly reduced under water deficit (Fig. 7b). In addition, the lowest values for shoot dry mass (Fig. 7a) and leaf area (Fig. 7c) were found in plants supplied with less NO_3_^−^ under low water availability.

**Figure 7.**
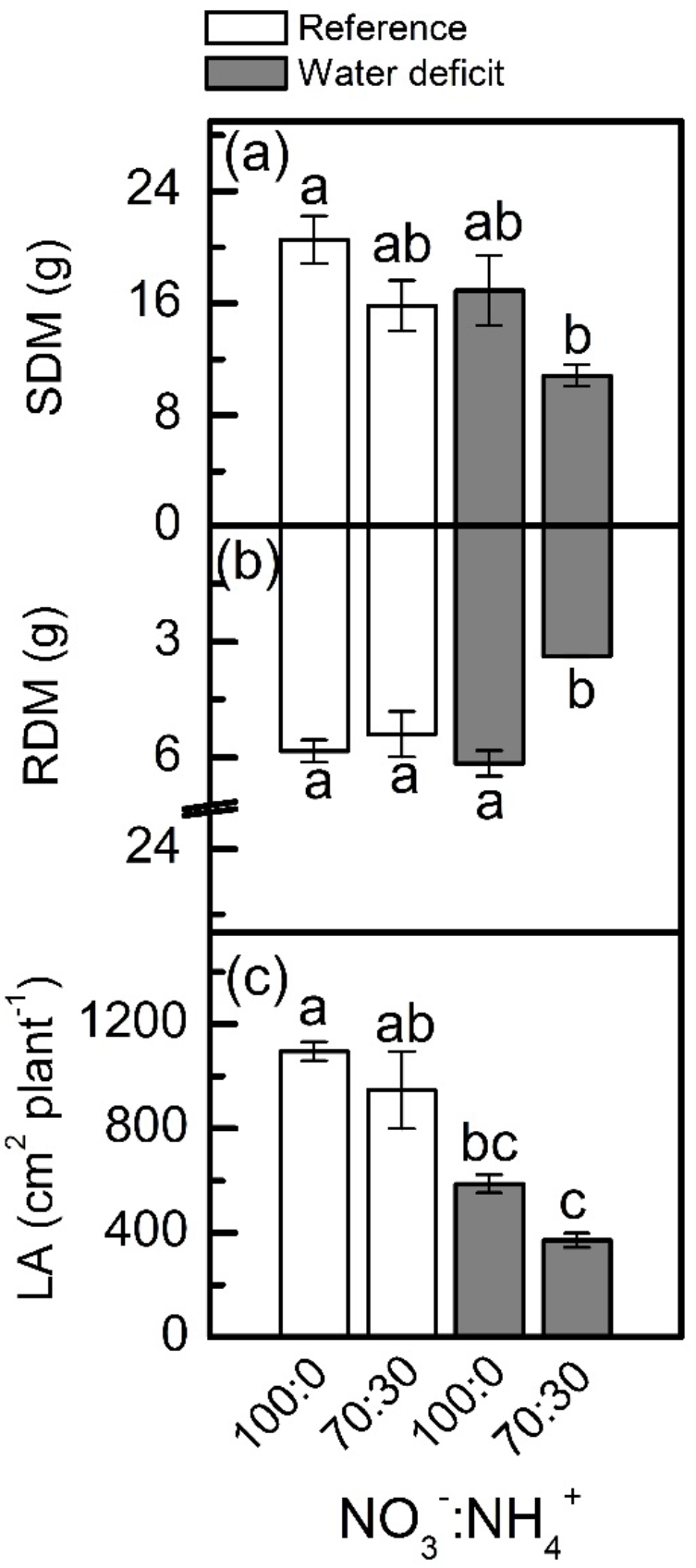
Shoot (SDM, in a) and root (RDM, in b) dry mass and leaf area (LA, in c) of sugarcane plants maintained well-hydrated (reference, white bars) or subjected to water deficit (gray bars) and supplied with varying NO_3_^−^:NH_4_^+^ ratios: 100:0 and 70:30. Bars represent the mean value of four replications ± se. Different letters indicate statistical difference among treatments (Tukey test, *p*<0.05).

#### Experiment II: Offsetting the benefits of NO synthesis induced by NO_3_^−^ supply

cPTIO – a NO scavenger – was sprayed on roots supplied with 100% NO_3_^−^ and facing water deficit. As consequence, the intracellular NO synthesis was reduced in leaves and roots (Fig. 8a,b) and plants showed lower photosynthetic rates and stomatal conductance under water deficit as compared to ones not sprayed with cPTIO (Fig. 9a,b). As found in experiment I, plants presented decreases in root dry mass due to water deficit when cPTIO was sprayed (Fig. 9d; Suppl. Fig. S2).

**Figure 8.**
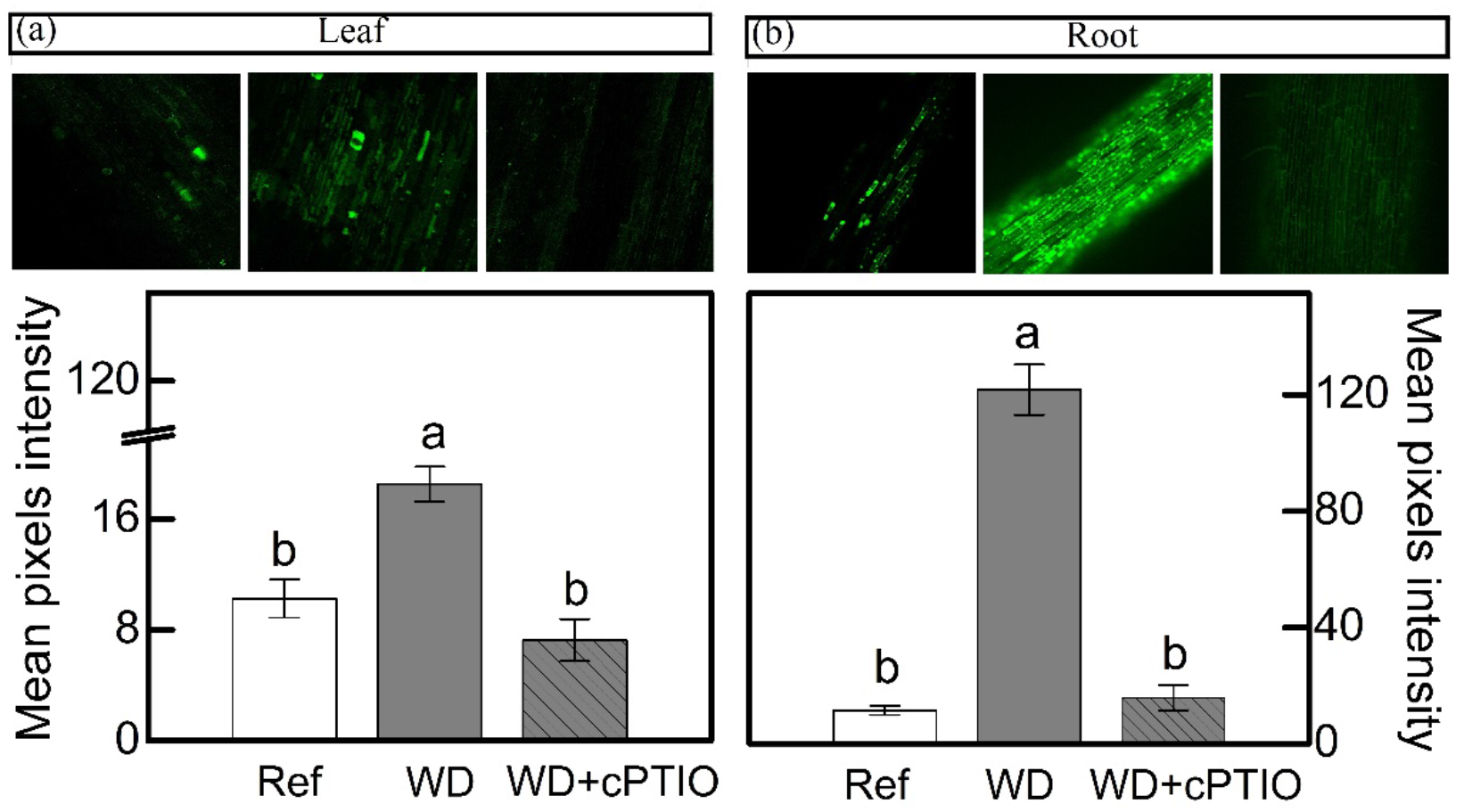
Confocal microscopy images showing intracellular NO synthesis in leaves (a) and roots (b) of sugarcane plants supplied with only NO_3_^−^ (100:0 NO_3_^−^:NH_4_^+^) and maintained well-hydrated (reference, white bars), subjected to water deficit (WD, gray bars) and subjected to water deficit and sprayed with cPTIO (WD+cPTIO, gray striped bars). Mean pixel intensities are also shown. Bars represent the mean value of four replications ± se. Different letters indicate statistical difference among treatments (Tukey test, *p*<0.05).

## Discussion

### Nitrate supply stimulates root NO production, improving photosynthesis and antioxidant metabolism of sugarcane under water deficit

Our findings revealed that nitrate reductase is an important enzymatic pathway for NO synthesis and also that sugarcane plants supplied with 100% NO_3_^−^ presented enhancement of drought tolerance. Here, we found higher NO_3_^−^ accumulation in roots under water deficit and receiving only NO_3_^−^ as source of nitrogen (Fig. 2b), which caused higher NO_2_^−^ production when compared to roots exposed to 70% NO_3_^−^ and 30% NH_4_^+^ (Fig. 2b,d). Such findings are supported by higher root nitrate reductase activity (Fig. 3b), which reduces NO_3_^−^ to NO_2_^−^ during the N assimilation pathway (Heidari *et al.*, 2011). As an alternative reaction, nitrate reductase may also reduce NO_3_^−^ to NO (Fancy *et al.*, 2016). In fact, the highest NO synthesis was found in roots under water deficit and supplied with only NO_3_^−^ (Fig. 6b) and it is known that NO_3_^−^ and NO_2_^−^ play a key role in NO synthesis through nitrate reductase (Vanin *et al.*, 2004; Yamasaki, 2005; Sun *et al.*, 2015). In *Physcomitrella patens,* low nitrate reductase activity was associated with drastic reductions in NO synthesis, further evidence that nitrate reductase is an important pathway for NO production in plants (Andrés *et al.*, 2015). It is worth noting that NO synthesis is low under non-limiting conditions, even in plants supplied with only NO_3_^−^ (Fig. 6). In general, increases in NO synthesis are expected under stressful conditions, when NO_2_^−^ accumulation occurs (Mur *et al.*, 2012).

In the last decades, rapidly increasing evidence has indicated NO as an important player in plant responses to environmental constraining conditions by inducing the antioxidant defenses (Hatamzadeh *et al.*, 2015; Silveira *et al.*, 2017b). During cell detoxification, O_2_^·−^ is dismuted to H_2_O_2_ by superoxide dismutase, which is rapidly eliminated by catalase and ascorbate peroxidase, producing H_2_O and O_2_ (Lázaro *et al.*, 2013). Here, we observed higher superoxide dismutase activity in roots under water deficit and supplied with 100% NO_3_^−^ (Fig. 5b), with root [O_2_^·−^] remaining similar among treatments (Fig. 4b). Interestingly, there was lower O_2_^·−^ accumulation in leaves under water deficit and supplied with only NO_3_^−^ (Fig. 4a), even with superoxide dismutase showing similar activity to the one found in plants supplied with 70% NO_3_^−^ and 30% NH_4_^+^ (Fig. 5a). As a possible explanation, such low leaf [O_2_^·−^] may be related to the interaction of this radical with NO, which generates peroxynitrite (ONOO^−^) and adds a nitro group to tyrosine residues – a process known as tyrosine nitration (Begara-Morales *et al.*, 2014; Wullf *et al.*, 2009). Although tyrosine nitration was originally considered as indicative of stress conditions, recent evidence suggests its role in cell signaling (Mengel *et al.*, 2013).

Root [H_2_O_2_] was lower in plants under water deficit that received 100% NO_3_^−^ as compared to ones supplied with 70% NO_3_^−^ and 30% NH_4_^+^ (Fig. 4d), indicating an efficient detoxification through increased root ascorbate peroxidase activity (Fig. 5d). In fact, the activation of antioxidant mechanisms to maintain ROS homeostasis often involves NO (Hatamzadeh *et al.*, 2015; Silveira *et al.*, 2015). Many reports show that exogenous NO improves abiotic stress tolerance, causing decreases in [H_2_O_2_] and lipid peroxidation (Gross *et al.*, 2013). Exogenous NO supply inhibits ROS accumulation in many plant species under stress conditions (Verma *et al.*, 2013), such as cucumber and rice under drought (Farooq *et al.*, 2009). Sugarcane plants supplied with GSNO – a NO donor – showed increases in the activity of antioxidant enzymes, such as superoxide dismutase in leaves and catalase in roots under water deficit (Silveira *et al.*, 2017b). In addition, the S-nitrosylation has a role in mediating the interplay between NO and other reactive signaling mechanisms, such as those involving ROS. For instance, S-nitrosylation of RBOHD causes its inactivation and thus reduces ROS formation through this pathway (Yu *et al.*, 2012). Such findings revealed that NO has an important role in controlling endogenous ROS levels.

Higher superoxide dismutase and ascorbate peroxidase in roots facing water deficit and receiving only NO_3_^−^ (Fig. 5b,d) may be a consequence of S-nitrosylation. In pea (*Pisum sativum*), S-nitrosylation increased the activity of cytosolic ascorbate peroxidase (Begara-Morales *et al.*, 2014). However, we noticed higher levels of S-nitrosothiols (SNOs) in roots under water deficit and supplied with NO_3_^−^ and NH_4_^+^ (Fig. 3f). At this point, one should consider that NO-mediated post-translational modifications on target proteins may be positive or negative (Nabi *et al.*, 2019). Some of these modifications may alter signaling pathways mediated by other ROS (Holzmeister *et al.*, 2014). According to Clark *et al.* (2000), S-nitrosylation can inhibit catalase activity, which implies that low level of S-nitrosylation can increase catalase activity during stress conditions, thus increasing ROS detoxification. In this way, higher [SNO] found in plants that received less nitrate (Fig. 3e,f) is associated with changes in the antioxidant system that lead to increases in leaf [O_2_^·−^] and root [H_2_O_2_] (Fig. 4a,d). It has been proposed that S-nitrosylation can regulate [H_2_O_2_] in plants, controlling both the antioxidant defense system and the ROS-producing enzymes (Ortega-Galisteo *et al.*, 2012; Yu *et al*., 2012).

Here, we found low accumulation of SNOs and high GSNOR activity in roots under water deficit that received 100% NO_3_^−^ (Fig. 3f,d). GSNOR can break down GSNO – a SNO, reducing GSNO levels and consequently decreasing the total cellular level of S-nitrosylation (Feechan *et al.*, 2005). Thus, it indirectly controls the overall SNOs within cells (Feechan *et al.*, 2005), suggesting that GSNOR may be crucial in regulating the cellular SNO pool. In fact, increases in GSNOR activity contributed to the reduction of S-nitrosylation in pea plants under salt stress (Camejo *et al.*, 2013). As GSNO is an NO donor, we can argue that increases in root GSNOR activity under water deficit and supplied with only NO_3_^−^ (Fig. 3d) are related to the reduction of GSNO levels and linked to high NO synthesis in roots (Fig. 6b). High levels of reactive nitrogen species (RNS) may be harmful to plants (Nabi *et al.*, 2019) and the absence of GSNOR activity in plants results in a significant increase in levels of SNOs and impairment of plant immunity (Feechan *et al.*, 2005), plant growth and development (Kwon *et al.*, 2012). Gong *et al.* (2015) demonstrated that absence of GSNOR activity increased the sensitivity of *Solanum lycopersicum* to alkaline stress due to the excessive accumulation of NO and SNOs, causing higher levels of endogenous S-nitrosylation and turning stomata insensitive to ABA.

Stomatal closure is the primary response of plants to water deficit, reducing the CO_2_ supply for photosynthesis and then decreasing biomass production (Machado *et al.*, 2009; Ribeiro *et al.*, 2013). Although water deficit had reduced the stomatal conductance, higher NO_3_^−^ supply alleviated such negative effects (Fig. 1b). Due to higher stomatal conductance, sugarcane plants supplied with 100% NO_3_^−^ showed an improvement in photosynthesis under water deficit (Fig. 1a). By integrating CO_2_ assimilation throughout the experimental period, plants supplied with only NO_3_^−^ fixed about 1.5 times more carbon than those supplied with NO_3_^−^ and NH_4_^+^. Such a response was also related to improvement of primary photochemistry, with plants showing higher conversion of light energy into chemical energy at the PSII level (Fig. 1c).

Under water deficit, plants supplied with 70% NO_3_^−^ and 30% NH_4_^+^ presented reduced root biomass as compared to those supplied with 100% NO_3_^−^, which were not affected by low water availability (Fig. 7b). Such increase in root growth was associated with higher NO content (Fig. 6b), as found by Silveira *et al.* (2016). At maximum water deficit, high NO synthesis was found in the root meristematic zone of plants supplied with 100% NO_3_^−^ (Suppl. Fig. S3). Several reports indicate that NO is involved in the regulation of root growth and developmental processes (Correa-Aragunde *et al.*, 2004; Lombardo and Lamattina, 2012; Sun *et al.*, 2015). The root system is able to perceive low water availability and to produce chemical signals that regulate the water flow from roots to shoots. NO is one of those chemical signals that stimulates root expansion and development (Xu *et al.*, 2017; Silveira *et al.*, 2016). Given the effects of NO on root growth, it is reasonable to assume a potential influence of NO mediating auxin signaling in roots. Correa-Aragunde *et al.* (2006) demonstrated that auxin-dependent cell cycle gene regulation was dependent on NO during lateral root formation in tomato plants. NO also modulates the auxin response during adventitious root formation in cucumber plants (Pagnussat *et al.*, 2002) and *Arabdopsis thaliana* (Lombardo *et al.*, 2006).

Overall, increases in NO content can trigger root development and improve water uptake, reducing the impact of low water availability on leaf water status and allowing higher stomatal conductance and photosynthesis, as noticed herein and also by Silveira *et al.* (2017). The novelty here is that we were able to induce NO synthesis in sugarcane plants by changing the nitrogen source. Such a finding has a practical consequence for sugarcane in the field as endogenous NO synthesis can be stimulated by increasing NO_3_^−^ supply. Apart from economic issues, our data give insights on how stress tolerance can be managed by common practices in agricultural systems and further development on this technique should be carried out with field-grown plants, where interactions among nutrients, soil-root interactions and soil nitrogen dynamics determine plant performance.

### Is sugarcane performance under water deficit really improved by NO?

Herein, we used 2-(4-carboxyphenyl)-4,4,5,5-tetramethylimidazoline-1-oxyl-3-oxide (cPTIO) – an endogenous NO scavenger (Akaike and Maeda, 1996) – to check if benefits induced by increasing NO_3_^−^ supply were related to NO. cPTIO drastically reduced the DAF-2DA in plants under water deficit, indicating lower NO accumulation in both leaves and roots (Fig. 8a,b). As consequence, plants showed even lower stomatal conductance and photosynthesis when compared to plants under water deficit and not supplied with cPTIO (Fig. 9a,b). cPTIO sprays also reduced root growth (Fig. 9d), as found previously (Fig. 7b). Taken together, our data clearly show that the improved performance of sugarcane plants supplied with only NO_3_^−^ were due to stimulation of NO synthesis under water deficit.

**Figure 9.**
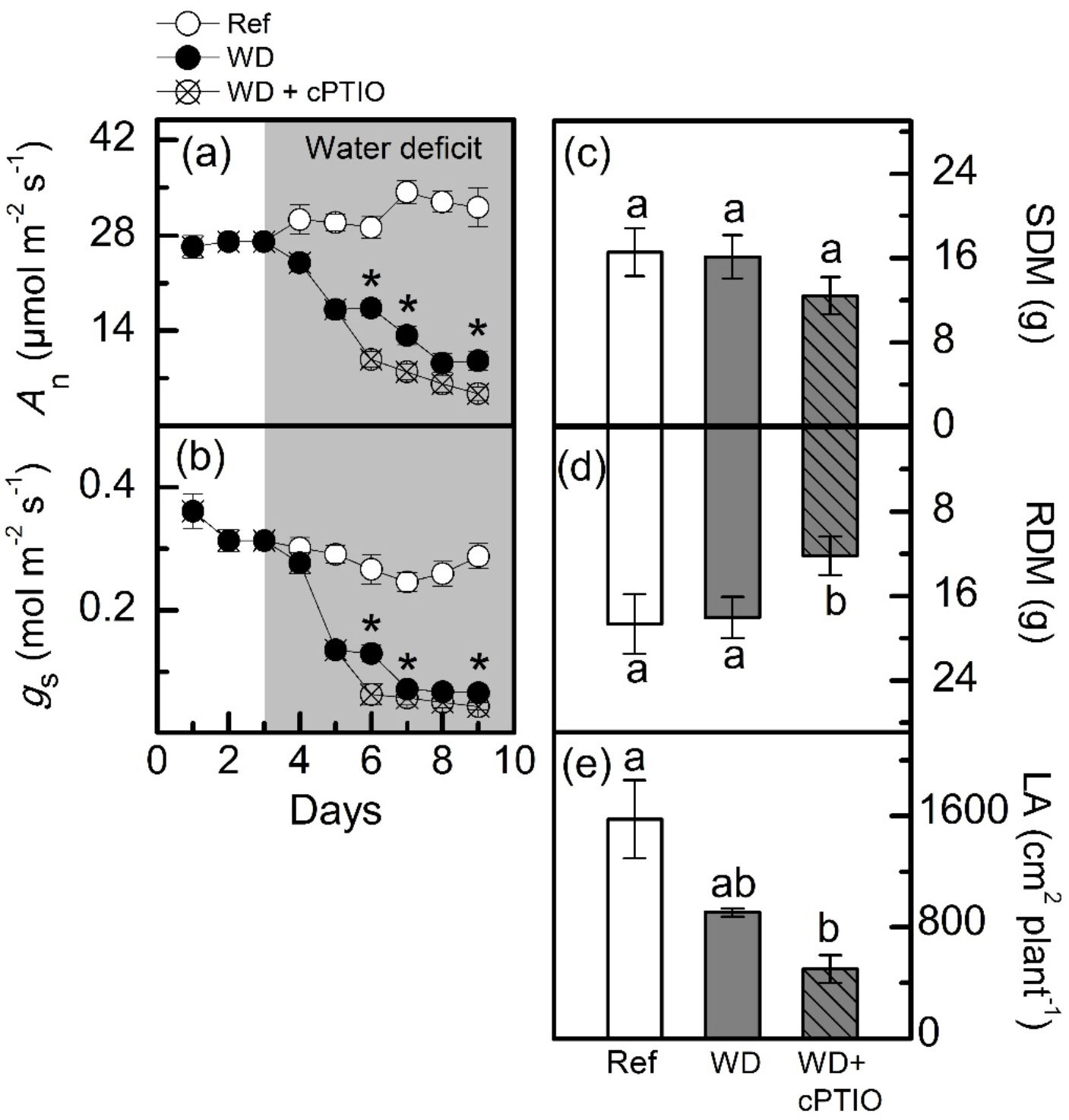
Leaf CO_2_ assimilation (*A*_n_, in a), stomatal conductance (*g*_S_, in b), shoot (SDM, in c) and root (RDM, in d) dry mass and leaf area (LA, in e) of sugarcane plants supplied with only NO_3_^−^ (100:0 NO_3_^−^:NH_4_^+^) and maintained well-hydrated (reference, white symbols and bars), subjected to water deficit (WD, black symbols and gray bars) and subjected to water deficit and sprayed with cPTIO (WD+cPTIO, crossed symbols and gray striped bars). Asterisks indicate significant differences between treatments under water deficit (a and b) and different letters indicate statistical difference among treatments (c-e) by the Tukey test (*p*<0.05).

## Conclusion

Sugarcane plants grown in nutrient solution containing only NO_3_^−^ as nitrogen source were more tolerant to water deficit and this response was associated with increased NO production and high nitrate reductase activity in roots. Herein, increasing NO_3_^−^ supply was enough to stimulate NO synthesis and alleviate the effects of water deficit on sugarcane plants by increasing the activity of antioxidant enzymes, photosynthesis, stomatal conductance and root growth. From a broad perspective, our data show that supplying more NO_3_^−^ during nitrogen fertilization may improve sugarcane tolerance and be beneficial to field-grown sugarcane.

## Supplementary data

**Fig. S1**. Activity and immunoblots of phosphoenolpyruvate carboxylase and ribulose-1,5-bisphosphate carboxylase/oxygenase in sugarcane plants under water deficit.

**Fig. S2**. Visual aspect of sugarcane plants under water deficit after NO scavenging through cPTIO spraying.

**Fig. S3.** Intracellular NO synthesis in apical sections of sugarcane roots.

## Acknowledgments

MDP and ABS acknowledge the scholarship provided by the São Paulo Research Foundation (FAPESP, Brazil; Grant numbers 2017/11279-7, 2018/08194-2). NMS acknowledges the fellowship granted by the National Program of Post-Doctorate (PNPD), Agency for the Specialization of Higher Education Personnel (Capes, Brazil). ECM, LS, ABS and RVR acknowledge the fellowships granted by the National Council for Scientific and Technological Development (CNPq, Brazil). MTP acknowledges the PhD fellowship granted by the São Paulo Research Foundation (FAPESP, Brazil; Grant number 201705029-8). ABS acknowledges the grant provided by the National Council for Scientific and Technological Development (CNPq, Brazil; Grant number 404815/2018-9). The authors are also very grateful to the National Institute of Science and Technology of Photonics applied to Cell Biology (INFABIC) for guiding our analyses with the confocal microscope.

